# Evidence that Dogs Can Use Temporal Difference in Odorant Arrival to Discriminate Odorant Mixtures

**DOI:** 10.64898/2026.06.26.734637

**Authors:** Iwan Downie, Paul Szyszka, Nathaniel J. Hall, Timothy L. Edwards

## Abstract

In turbulent environments, odorants from different sources arrive at different times, potentially providing cues for odor source segregation. In several invertebrate species, short differences in odorant onset enable freely moving animals to discriminate odorant mixtures. In vertebrates, however, studies of sensitivity to odorant onset asynchrony have been conducted under highly constrained sampling conditions, such as with odor delivery tightly coupled to respiration. In this study, we investigated whether domestic dogs could detect odorant onset asynchrony in odorant mixtures under conditions that preserve key features of natural odor sampling. Dogs performed a discrimination task in which odor stimuli were presented as ongoing pulse trains that began independently of animal behavior, avoiding artificial synchronization of odor delivery with sniff cycles. Dogs were trained to discriminate between mixtures of two odorants with synchronous onsets and mixtures with asynchronous onsets. Of the dogs trained, one was able to discriminate odorant onset asynchronies as short as 633 ms. Dogs also displayed sensitivity to auditory stimulus onset asynchrony, discriminating auditory asynchronies as short as 30 ms. These results provide the first demonstration of temporal sensitivity in canine olfaction and the first evidence that vertebrates can use odorant onset asynchrony under conditions that permit free odor sampling.

## Introduction

Natural odor plumes consist of intermittent odor filaments that generate fluctuating temporal patterns of odorant arrival at the olfactory organs (Celani et al., 2014; Crimaldi & Koseff, 2001; Stark et al., 2026). These temporal patterns provide spatial cues that animals could use to locate and track odor sources (Jayaram et al., 2022; Marin et al., 2021; Michaelis et al., 2020; Nag & Van Breugel, 2024; Szyszka et al., 2023).

Domestic dogs (*Canis familiaris*) have been bred for millennia for their many functions, including olfaction (Marshall-Pescini & Kaminski, 2014). Today, dogs’ olfactory abilities have been employed in various roles (Hall et al., 2016; Kokocińska-Kusiak et al., 2021), including policing and security (Vyplelová et al., 2014), and conservation (Grimm-Seyfarth et al., 2021). Although dogs show high sensitivity to odor identity and concentrations and are highly effective at detecting target odors in complex environments (Kokocińska-Kusiak et al., 2021; Moser et al., 2019), it remains unclear whether they use temporal odor cues to segregate overlapping odor plumes originating from different sources. Such segregation of target odors from background odors and other odor sources is a fundamental challenge in olfaction (Linster et al., 2007; Riffel et al., 2014; Rokni et al., 2014; Stevenson & Wilson, 2007).

For example, when a dog searches for a missing person in an environment where other people have recently passed through, it needs to discriminate between the odorants emitted by the target from those emitted from other people whose odorants may have mixed in the air.

One proposed mechanism for solving this odor segregation challenge relies on temporal stimulus cues, particularly stimulus onset asynchrony: differences in the timing of odorant arrival between odor plumes from different sources (J. J. Hopfield, 1991). In audition, stimulus onset asynchrony contributes to source segregation, as illustrated by the cocktail party effect, in which sounds are grouped based on their relative timing (Carlyon, 2004). Analogously, temporal differences in odorant arrival may enable animals to segregate odor sources (Ackels, 2022; J. J. Hopfield, 1991; Szyszka et al., 2023).

Stimulus onset asynchronies between odorant encounters originating from different sources are expected to span a broad range, from milliseconds to seconds, depending on source separation, flow structure, and distance from the source (Celani et al., 2014; Crimaldi & Koseff, 2001; Soltys & Crimaldi, 2015; Stark et al., 2026). Studies with insects (Andersson et al., 2011; Baker et al., 1998; Saha et al., 2013; Sehdev et al., 2019; Sehdev & Szyszka, 2019; Szyszka et al., 2012), crustaceans (Weissburg et al., 2012), and slugs (J. F. Hopfield, 1989) provide evidence that animals can exploit stimulus onset asynchronies spanning from milliseconds to seconds to segregate odor sources.

Recent studies indicate that vertebrates are also sensitive to odorant onset asynchrony, albeit in the tens of milliseconds range. Mice successfully discriminate between odorant mixtures where odorants were presented either fully synchronously (AB) or asynchronously (rapidly fluctuating A-B stimuli), with odorant onset asynchrony as low as 25 ms (Ackels et al., 2021). Humans can discriminate between the order of odorant stimuli (A-B vs. B-A), with different asynchrony thresholds reported alongside different study methodologies. Wu et al. (2024) showed that humans can discriminate between odor mixtures that differ only in the relative timing of their components, distinguishing A→B from B→A sequences that differed by 120 ms in relative timing. Ding et al. (2019), Laing et al., (1994), and Perl et al. (2020) used an approach in which air was delivered near the participants’ noses; they were instructed to inhale at a time that corresponded with odorant mixture delivery. Without any instructions on the onset asynchrony in the stimuli, Perl et al.’s participants were unable to distinguish between synchronous and asynchronous mixtures, even at 600 ms asynchrony values.

However, when instructed about the ordering of the stimuli, they were able to discriminate with asynchrony as little as 300 ms. Instead of a “same or different” task, Ding et al. asked participants which odorant arrived first and found they could do so when the component odorant onset was separated by as little as 450 ms. Similarly, Laing et al. found that, when asked which odorant arrived first, participants could do so when onset was separated by between 92 ms and 312 ms, depending on the odorant mixture used.

Importantly, in all of the previous studies with vertebrates, the nature of the stimuli or the manner in which they were presented to the organism lack the complexity of natural odor stimuli or the way that organisms typically interact with them. In the human studies, the odorant stimuli were always synchronized with the sniff cycle, and participants were exposed to a single odorant onset in each trial. Ackels et al. (2021) presented A-B sequences but with no overlap in components during the asynchronous stimuli; this separation between components does not occur in natural odor plumes. Although these studies provide valuable information about the fundamental sensitivity of these olfactory systems to odorant onset asynchrony, they do not establish whether vertebrates can detect such temporal onset asynchronies under complex naturalistic conditions.

In the present study, we investigated whether dogs are sensitive to odorant onset asynchrony using an experimental design that preserves key features of natural odor sampling. Odorants were presented as continuous pulse trains that began independently of the dogs’ behavior and persisted until a response occurred, allowing dogs to sniff freely and sample the odor plume in a manner more closely resembling natural conditions than previous studies. Stimuli were generated using an apparatus, adapted from Raiser et al. (2017) for dogs, which provided precise control over stimulus onset asynchrony without directly coupling odorant delivery to respiration. After establishing that dogs could perform odor discrimination tasks using odorant onset asynchrony, we quantified a dog’s threshold sensitivity to odorant onset asynchrony using overlapping odorant pulse trains that approximate the complexity of natural plume structures.

## Methods

### Participants

Participants were recruited from the community in Waikato through posters and social media. Dogs were initially screened for food motivation, aggressive tendencies, and separation anxiety via questionnaire. Dogs that met criteria were assessed in person for their response to food, the experiment space, and the researchers. A total of eight dogs met initial criteria to begin training, with five being withdrawn over the course of training before completing final testing (Table 1). Reasons for withdrawal were predominantly due to dogs failing to engage with the task sufficiently (Bojji, Susie, Ellie).

**Table 1.**
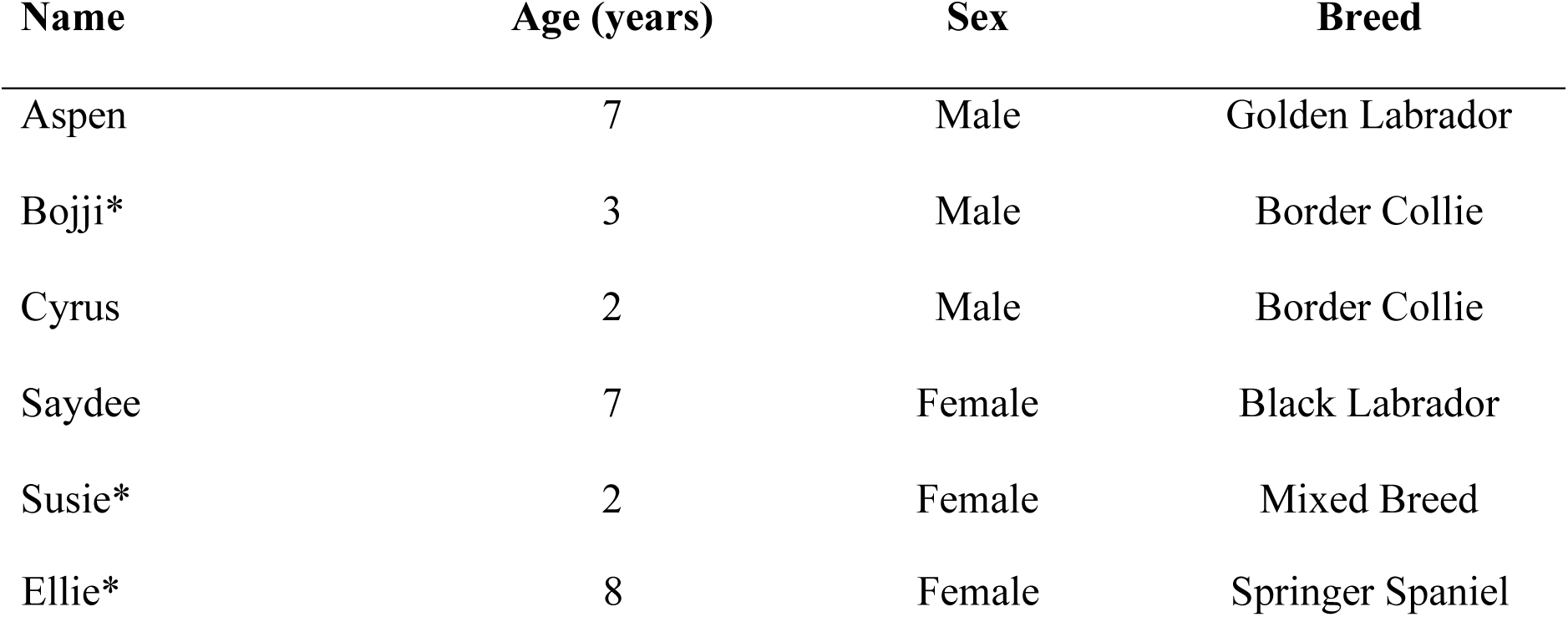
Participant Information. All participants were desexed. *Withdrawn.

Participants lived with their owners for the duration of the study, attending training sessions at the University of Waikato’s Scent Detection Research Group facility, during which they were cared for by researchers. All experimental procedures were approved by the University of Waikato Animal Ethics Committee (protocol 1209).

### Apparatus and Setting

As part of this study, an automatic apparatus was developed which could deliver odorants with control over odorant arrival timeat a sample port where dogs could sample odorant mixtures and respond to them. The apparatus was developed using Raiser et al.’s (2017) fast olfactory stimulator as the apparatus core, with the rest of the apparatus developed for dogs (Figure 1).

**Figure 1.**
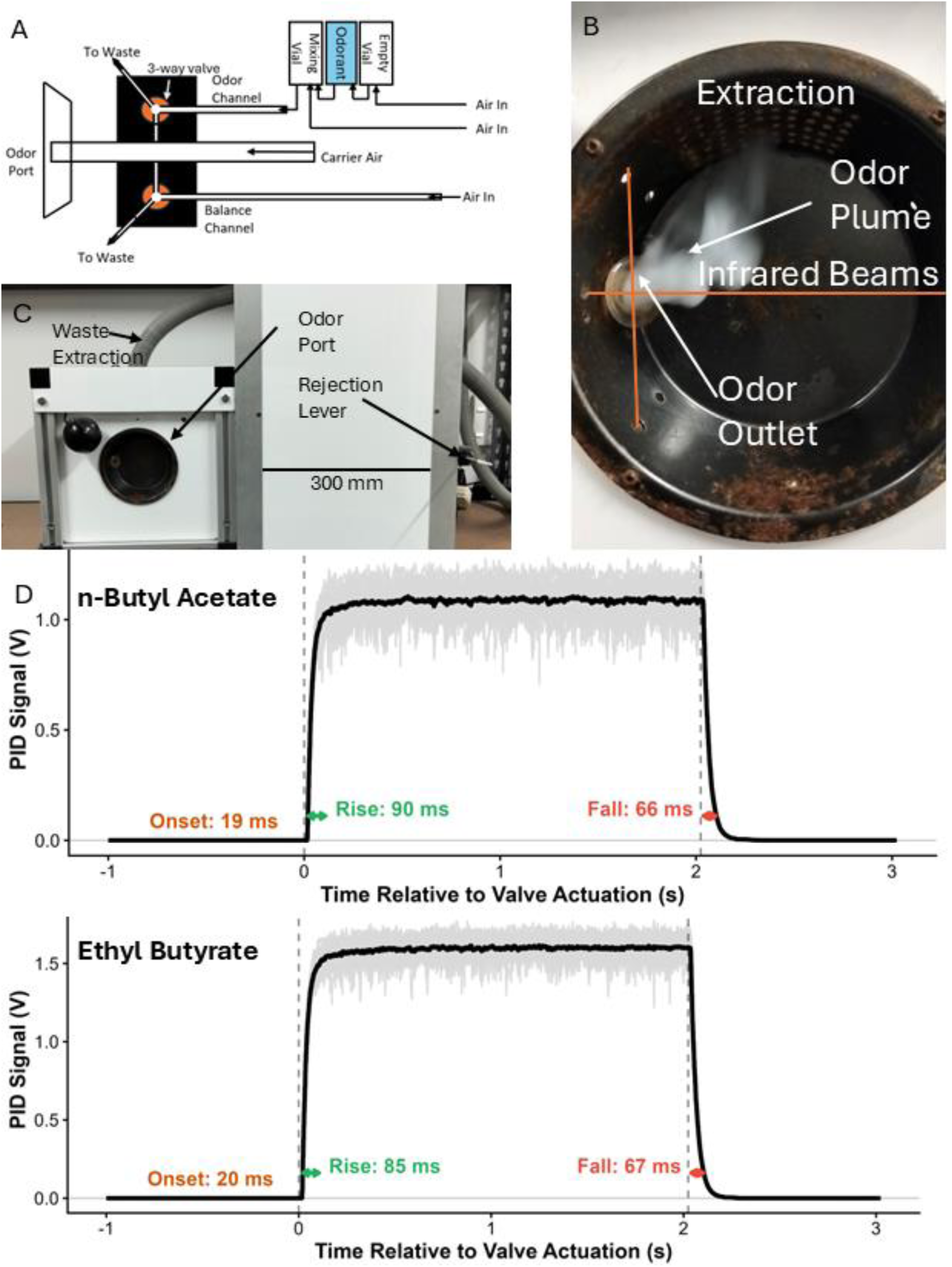
Dog-adapted apparatus for controlled odorant onset asynchrony. (A)Scheme of one odorant channel of the stimulator; this layout was replicated to allow 2 odorant channels. Odorant pulses are presented by diverting air flow in the odorant channel from waste into carrier air through the three-way valve and simultaneously air flow in the balance channel is diverted from the carrier air to the waste. This ensures that the total air flow volume at the odor port remains constant. (B) and (C) show the apparatus in its experimental configuration. In B, the white vapor was used to visualize airflow across the port from the outlet (port 100 mm diameter × 50 mm deep, outlet 9mm inner diameter). Vapor was not present during dog experiments and was used only for verification of airflow. The infrared beams were used to monitor when the dog’s nose was in position over the odor outlet, the tube connecting the odor port to the apparatus core. (D) Photoionization detector (PID) signal showing the mean of 35, 2 s-long odorant pulses of n-butyl acetate (Sigma-Aldrich, 99.5% purity), and 37, 2 s-long odorant pulses of ethyl butyrate (Acrōs Organics, 99% purity) in black. Output voltage signals from the photoionization detector (PID) were sampled at 1000 Hz. Each pulse signal was adjusted by subtracting the mean voltage of the last 500 ms before valve actuation to account for sensor drift across subsequent recordings. PID measurements were taken from 2 mm outside the odor outlet, in the approximate location dogs were required to sample. The grey lines represent individual pulses overlayed to display concentration variability across pulses.

The apparatus consisted of a polyether ether ketone (PEEK) core using high speed three-way valves (LFAA1208010H Lee Manifold Mount Valve, Lee) to control odorant release into a carrier airstream. Air was fed from a compressor through a carbon filter (APF23CA, KSI) and a water filter (APF23SMA, KSI) to remove contaminants and water vapor before being distributed into two odorant channels, two balance channels, and one carrier channel. Waste air was collected from all valves and the odor port and vented into an exhaust system. Pure odorants were contained in 7 ml glass vials (diameter = 10mm, height = 40mm) positioned in-line between the odorant channel air inlet and the apparatus core (Figure 1A). Clean air was continuously fed through the headspace of the vials at a rate of 0.1 L/min to ensure a steady-state concentration of odorants in the headspace due to a balance between odorant evaporation and removal by the air flush. As a result of this continuous airflow, the headspace odorant concentration never reached saturation. Odorant concentration was further diluted by mixing the sampled headspace with un-odorized air in a separate glass mixing vial with clean air added at a rate of 0.2 L/min. This odorized air was then mixed with the carrier stream (flow rate = 3.0 L/min) in the apparatus core, further diluting concentration. All airlines which encountered odorized air were either made from PTFE or replaced at any time odorants were changed to prevent cross contamination. Odorized air was fed through three-way valves with one inlet and either routed to the waste stream or into the carrier airstream (3 L/min). To maintain constant airflow through the apparatus, balance valves were supplied with air at the same flow rate as the odorant channels and fed air into the carrier at a flow rate of 0.3 L/min when no odor stimulus was presented.

The change of odorant concentration after valve actuation was measured using a photoionization detector (miniPID 200B, Aurora Scientific). We then calculated odorant onset, rise times, and fall times. Onset time was calculated by identifying the peak amplitude of each PID signal and then searching backwards from the peak to identify the first time point at which the signal reached 5% of the peak amplitude. The mean onset time was 19.03 ms (SD = 1.56 ms) for 35 n-butyl acetate pulses, and 20.22 ms (SD = 1.40 ms) for 37 ethyl butyrate pulses. Rise time was calculated as the time required for the PID signal to rise from 10% to 90% of its peak amplitude. The mean rise time was 89.03 ms (SD = 22.57 ms) for n-butyl acetate, and 84.81 ms (SD = 14.02ms) for ethyl butyrate. Fall time was calculated as the time for the PID signal to fall from 90% to 10% of its peak amplitude. The mean fall time was 65.54 ms (SD = 5.92 ms) for n-butyl acetate and 66.89 ms (SD = 4.37 ms) for ethyl butyrate.

The apparatus was located in a 3m × 6m room and controlled from an adjoining room using custom software. The apparatus core was contained in a 300mm high × 300mm wide × 400mm deep box, with the interaction points attached. Interaction points on the apparatus consisted of an odor port housing infra-red sensors to monitor dog interaction with the odor stimuli and an omnidirectional switch located 50cm to the right of the odor port (Figure 1C). Olfactory stimuli were delivered from the apparatus core to the odor port through an outlet (internal diameter = 9 mm). Experiments were monitored and recorded using three closed circuit cameras positioned around the experiment room. Reinforcement was provided in the form of kibble delivered by a Treat&Train® food dispenser or Possyum (a semi moist dog food) delivered using a carousel feeding system.

### Odorants

Pentyl acetate (Sigma-Aldrich, 99% purity) and 1-hexanol (Sigma-Aldrich, 98% purity) were used for initial testing as they are known to be discriminable by dogs; however, these were not used for timing related tasks due to slow predicted rise and fall times of the odorant concentration due to their relatively low vapor pressure. Specifically, due to the relatively short pulse lengths used, the distinction between the synchronous and asynchronous stimuli was not reliable, with both odorants potentially being present at similar concentrations throughout asynchronous stimuli. Therefore, for odorant onset asynchrony discrimination, we initially used ethanol (Sigma-Aldrich, 99.5% purity) and ethyl acetate (Sigma-Aldrich, 99.5% purity). However, due to some dogs exhibiting avoidance responses, these were changed to n-butyl acetate (Sigma-Aldrich, 99.5% purity) and ethyl butyrate (Acrōs Organics, 99% purity), which did not produce avoidance. These were chosen as they enabled fast rise and fall times when measured with a PID (Figure 1D) while having low enough volatility to be used at detectable concentrations with the flow rates required for fast odor onset. Odor stimuli were presented as pulse trains to approximate the intermittent structure of odor stimuli encountered in turbulent plumes, allowing dogs to sample odor stimuli across multiple sniff cycles rather than at a fixed phase of respiration.

### Training

Following acceptance into the project, dogs were habituated to the apparatus, researchers, and automated feeders. They were allowed to explore the experimental room freely, with the apparatus running and the feeder being triggered to dispense food reinforcement, until they approached the feeder within three seconds of food being dispensed.

Once the feeder had been conditioned as a reinforcer, it was used to train a nose-hold inside the sample port using the method of successive approximations to desired behavior (shaping). Dogs were shaped to hold their nose at the intersection of two infrared beams in front of the odor port, breaking both beams for 1,500 ms, which served as the indication response and was indicated by a tone from the apparatus. Activation of the omnidirectional switch was then shaped. The response topography was not constrained; any response that activated the switch was reinforced. The switch was programmed to advance to the next trial without reinforcement and activation of the switch by the dog was treated as a rejection response, provided the dog had first completed an investigation, defined as breaking the infrared beams for more than 500 ms. After completing habituation and shaping procedures, dogs progressed to odor discrimination training (Figure 2).

**Figure 2.**
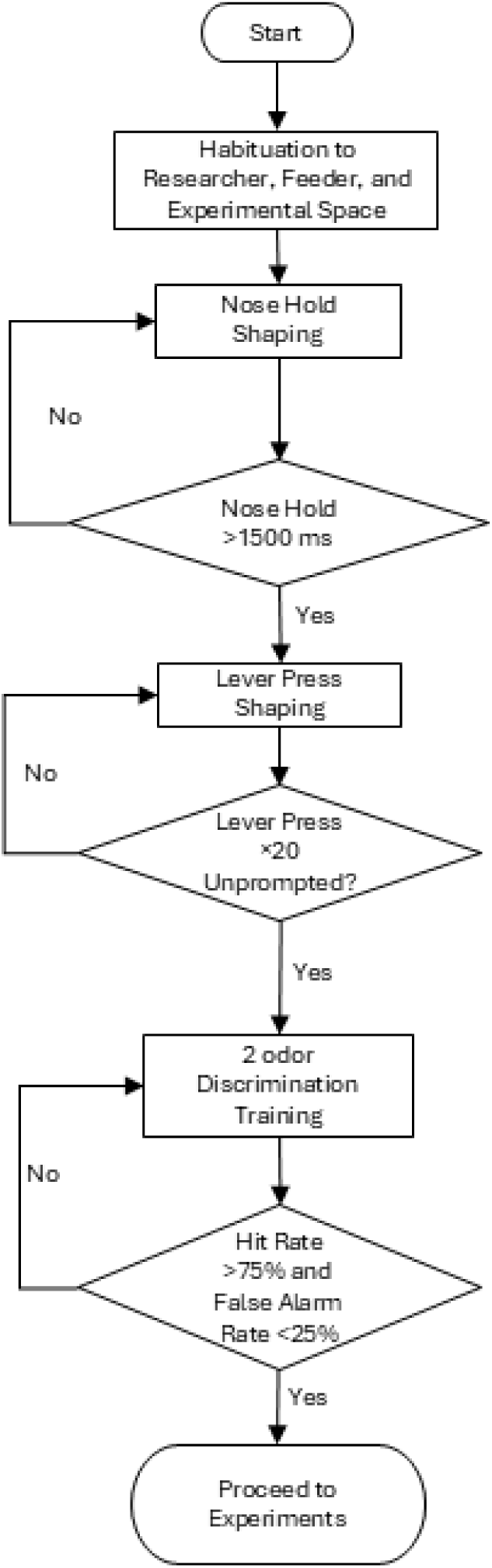
Sequence of habituation, shaping, and odor discrimination prior to onset asynchrony experiments 1 and 2

For odor discrimination, dogs were first trained through operant conditioning to discriminate between a single odorant A or B. This was done to ensure the dogs could reliably complete a scent detection task using the apparatus. During this stage of training, experimenters systematically removed themselves from the room to reduce opportunities for prompting. During discrimination training, a nose hold in the presence of the target (A) was reinforced using the automated feeder and recorded as a hit. A nose-hold in the presence of a non-target (B) was not reinforced and treated as a false alarm. To advance to the next trial during non-target trials, dogs had to perform a rejection by actuating the lever after completing an investigation. Investigations were defined as the dog placing their nose in the odor port, breaking the infrared beams for at least 500 ms. Any rejection response attempted before this time was not valid and would not advance the trial. This investigation ensured that dogs sampled from the odor port before completing a response behavior, which was a requirement for experiments involving odorant onset asynchrony to ensure exposure to odorant onsets. The rejection was not directly reinforced by food but triggered the start of the next trial in a session. The threshold for an indication response was varied individually until false alarm rates decreased, with all dogs ultimately working with a 5000-6000 ms indication threshold. This indication response requirement is a key component of the method used here, as when only the indication is reinforced the threshold must be high enough to ensure a high enough cost for false alarms. A more detailed description of this type of procedure and the influence of the indication threshold on response accuracy can be found in Edwards et al. (2022).

For initial training, pentyl acetate was used as the target, and 1-hexanol was used as the non-target. Before the end of training, dogs were also trained to discriminate between ethyl butyrate and n-butyl acetate to ensure they could discriminate between the individual odorants before odorant onset asynchrony was introduced. All sessions consisted of 19 trials throughout all experiments, with dogs completing between four and six sessions per day depending on each dog’s motivation to work, with at least a 10-minute-long break between sessions. Trials initially consisted of ten non-target trials and nine target trials, with the ratio later varied per dog as training progressed. The ratio of non-target trials was increased if a dog had greater than 90% hit rate and greater than 50% false alarm rate for at least 5 consecutive sessions. Once dogs performed this task at above 75% hit rate and below 25% false alarm rate for two out of three consecutive sessions, they were advanced to Experiment 1, which maintained the ratio between targets and non-targets identified during training.

### Experiment 1 - Estimating the Detection Threshold for Odorant Onset Asynchrony by Titration

The detection threshold for odorant onset asynchrony was first estimated using a discrimination task between a synchronous target and an asynchronous non-target. Nose holds were reinforced in the presence of a synchronous stimuli (AB), whilst all asynchronous (A-AB-B) stimuli were not reinforced and required a lever press to end the trial. Initially, dogs were trained using 500 ms pulses delivered at 1 Hz which were either fully synchronous target or fully asynchronous non-target (Figure 3). Once hit rate and false alarm rate were above 75% and below 25% respectively over three sessions, we advanced to threshold investigations where odorant onset asynchrony was reduced using titration. Throughout the titration procedure the ratio of target and non-target trials was held constant at the ratio identified for each dog during the training procedure.

**Figure 3.**
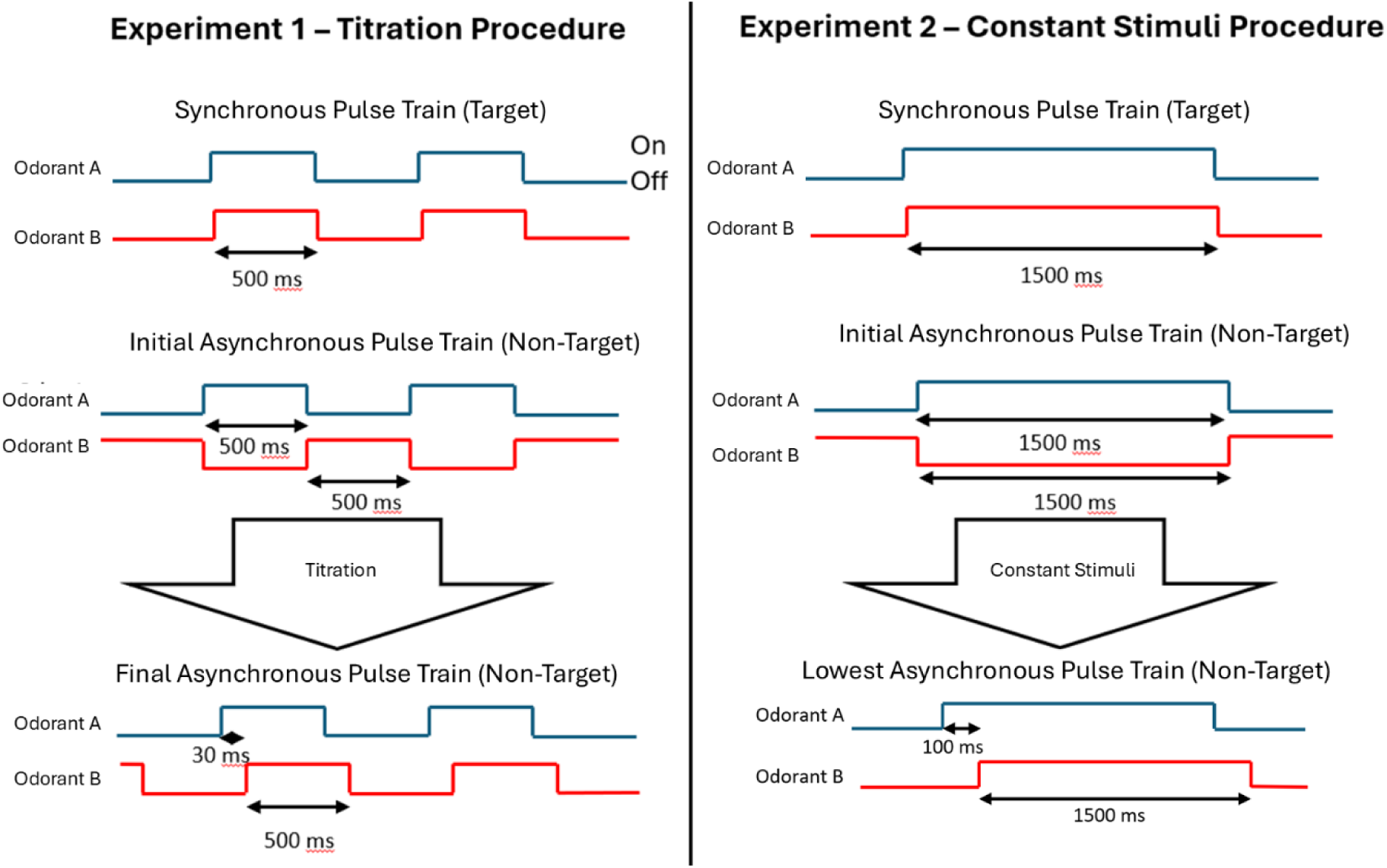
Odorant Onset patterns of binary odorant mixtures used for titration and method of constant stimuli experiments. Odorant Onset patterns of binary odorant mixtures used for threshold estimation in Experiment 1 (left) and 2 (right). The order of odorants A (ethyl butyrate) and B (ethyl acetate) was kept constant, with A always being presented first in the sequence. Reinforcement was always paired with the synchronous stimulus (AB).

During this titration experiment odorants were always presented for 500 ms per 1 second of the pulse train in each type of stimulus with the only change being to the asynchrony of onset and offset times. Onset asynchrony refers to the time between the valve for odorant A switching on and the valve for odorant B switching on.

Asynchrony was reduced by 100 ms if dogs achieved better than 75% hit rate and less than 25% false alarm rate for at least two out of three consecutive sessions. Asynchrony was reduced using these criteria until performance fell below 75% hit rate with a false alarm rate greater than 25% for 3 sessions, which did not have to be consecutive; or if the dog failed to meet criteria to reduce asynchrony within 10 sessions. Following this, asynchrony was increased by 100 ms and sessions were repeated until performance met the previous criteria. Following an increase in asynchrony, reduction increments were reduced to 10 ms steps. This process was repeated for all dogs until performance fell to chance levels for a given asynchrony value for 10 sessions after a second increase in odorant onset asynchrony.

After completing the titration procedure, dogs were returned to a level of asynchrony with which they had previously achieved greater than 75% hit rate and less than 25% false alarm rate. Odorants were then replaced with blank air to test for control of behavior by non-olfactory stimuli, where dogs were expected to perform at chance performance if their behavior was under the control of the olfactory stimuli. These tests indicated that dogs’ responses were indeed being controlled by non-olfactory stimuli. We therefore implemented a new auditory masking system and employed a new method of establishing odorant onset asynchrony thresholds in Experiment 2.

The auditory masking system used a multi-valve array which randomly activated five “dummy” valves (three-way valves identical to those used for the odor and balance channels) attached to the apparatus core with no air feed. The masking system was controlled by a microcontroller which randomly selected an actuation delay from an even distribution between 50-150 ms for each of the five valves. This delay was relative to the last actuation of the valve or the start of the trial, with each valve having a new delay selected following every actuation. Each valve was actuated following the randomly selected delay, and a new delay was selected. Following repeated evaluations in which dogs were tested with the modified masking both in the presence of odorants and under blank-air conditions, performance in the absence of non-olfactory stimuli fell to chance levels, indicating that discrimination was no longer supported by non-olfactory cues.

### Experiment 2 – Odorant Onset Asynchrony Threshold Estimation Using the Method of Constant Stimuli

To estimate the threshold for odorant onset asynchrony discrimination using olfactory cues, we applied a version of the method of constant stimuli (a psychophysical research technique), where all stimulus onset asynchronies to be evaluated were presented an equal number of times and randomly distributed across sessions using pulse trains as shown in Figure 3 (right). This constant stimulus method was adopted due to it requiring less time to complete than the titration method used in Experiment 1, which we felt may have contributed to the high withdrawal rate of dogs at that stage. Dogs were returned to the synchronous versus completely asynchronous condition and onset asynchrony was increased in 500 ms increments until an onset asynchrony was identified where false alarm rate was below 25% and hit rate was above 75% for at least two out of three consecutive sessions. The onset asynchrony required for above chance performance identified as 1,500 ms with 100 % asynchronous stimuli presented at 0.33 Hz. Once performance met target levels, dogs completed 24 sessions, each 19 trials long. In these sessions, non-target stimuli had variable onset asynchronies of 500 ms, 700 ms, 900 ms, 1,100 ms, 1,300 ms, and 1,500 ms. These were presented in a randomized order with all asynchronies being presented a total of 50 times. Throughout this phase, reinforced synchronous stimuli were included, with 6 synchronous trials and 13 asynchronous trials per session. Trial order within each session was determined using a computer-generated pseudorandom permutation based on a Mersenne Twister random number generator, ensuring equal probability of placement for all trials. *d’* was calculated as a measure of sensitivity for each level of asynchrony using Z-values for the number of hits and false alarms per session with the equation *d*′ = *Z*(*H*) − *Z*(*FA*).

Performance remained above chance for all onset asynchronies tested, meaning a threshold could not be identified and suggesting the range in asynchronies used was not sufficient to draw conclusions. Therefore, the process was repeated with 100 ms increments between 100 ms and 1,500 ms for 30 trials per increment to increase resolution in threshold estimates and to ensure the threshold fell in the range tested. After completing 50% of the planned test sessions, control probes were conducted in which odorants were replaced with blank air stimuli for three consecutive sessions to ensure discrimination was under control of olfactory stimuli before completing the remaining sessions.

## Statistical Analysis

To evaluate discrimination between target and non-target stimuli, sensitivity was calculated using *d’* for each tested onset asynchrony with chance represented by value ≤ 0 and a value ≥ 3 generally considered to be near perfect performance. *d’* was calculated from the z-scores for hit rate (*Z(H))* and false alarm (Z(FA)) proportions for the final three sessions completed at each odorant onset asynchrony during Experiment 1, and for all trials during Experiment 2. A log-linear correction was applied to avoid infinite *Z-scores*.

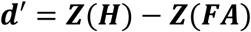

Finally, to find the estimated threshold for chance discrimination in Experiment 2, data were fitted to a four-parameter logistic sigmoidal model using the l.4 model in the drc package in R (V4.5.2). The function approximates a cumulative Gaussian model, modelling sensitivity as a function of onset asynchrony. The curve corresponds to the function:

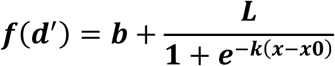

Where *b* represents the minimum *d’* recorded, *L* represents the range in recorded *d’*, and *x* is the onset asynchrony. The inflection point of the fitted curve was taken as the perceptual threshold. Uncertainty in the model fit was characterized by calculating a 95% confidence interval.

## Results

### Experiment 1 – Titration-Based Threshold Estimation and Identification of Non-Olfactory Cues)

Following training, two dogs (Aspen and Saydee) completed all stages of the titration protocol. Both dogs performed with greater than chance performance at onset asynchronies as low as 30 ms (Figure 4). In control sessions (with no odorants present), Aspen and Saydee continued to discriminate between synchronous and asynchronous stimuli with significantly above chance performance. This indicates that dogs relied on auditory cues rather than olfactory cues to complete the task, suggesting auditory sensitivity to onset asynchrony at least as low as 30 ms. Cyrus achieved greater than chance performance with onset asynchrony as low as 200 ms, but testing was stopped following the identification of other dogs using auditory cues to complete the discrimination task.

**Figure 4.**
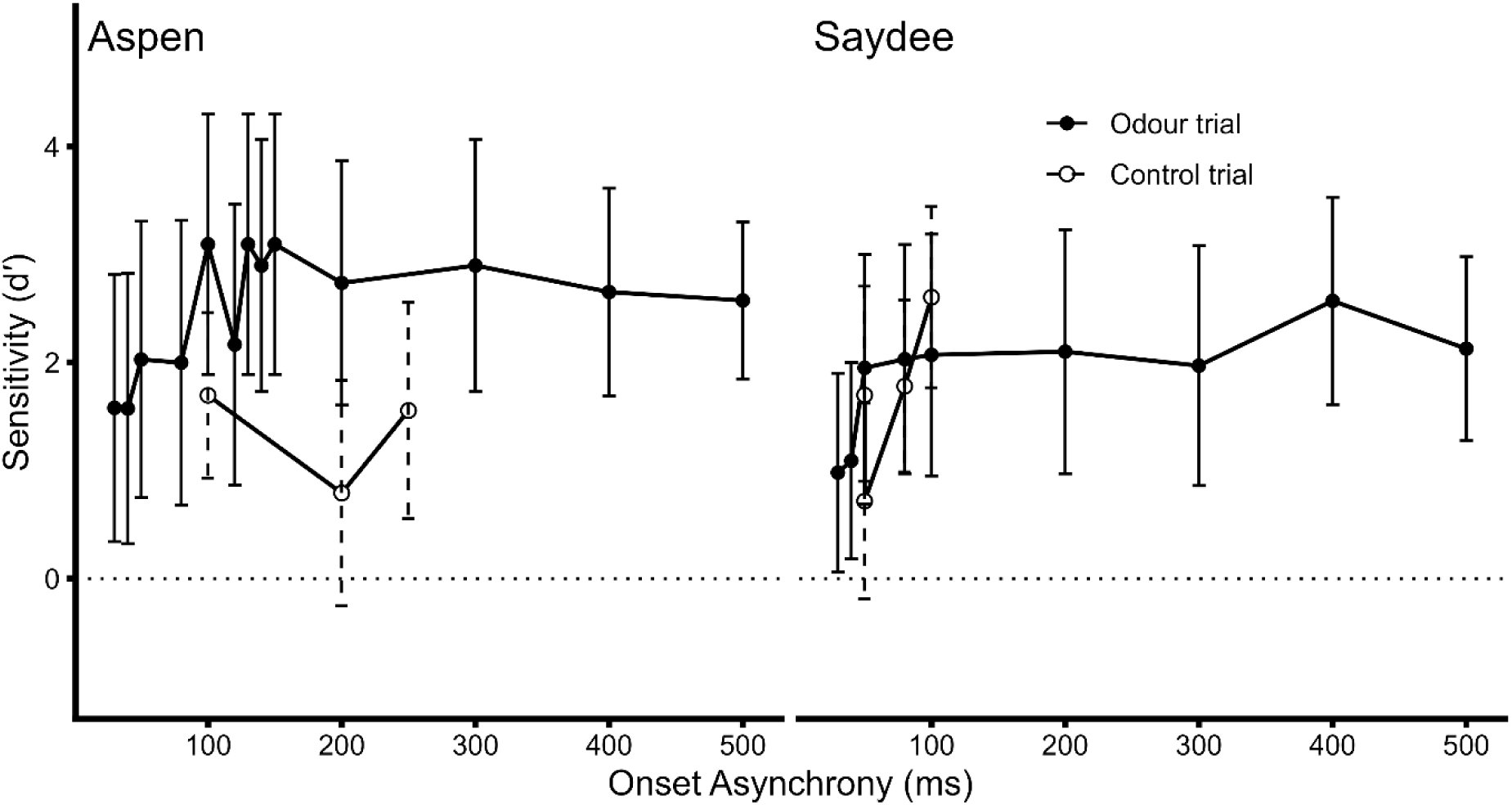
Discrimination between synchronous and asynchronous stimuli using auditory cues with a stimulus onset asynchrony threshold of 30 ms or below. Error bars represent 95% confidence intervals for calculated d’ values. Onset asynchrony values below 100 ms consisted of 80 ms, 50 ms, 40ms, and 30 ms. Aspen and Saydee performed significantly above chance with onset asynchrony as low as 30 ms.

### Experiment 2 – Constant-Stimuli Estimation of Odorant Onset Asynchrony Thresholds

Once behavior was reliably brought under olfactory stimulus control and the use of auditory cues was excluded, all dogs completed discrimination training with onset asynchronies of 1500ms presented at 0.33 Hz. At this stage, Saydee and Cyrus did not complete sufficient sessions due to motivation issues, leaving Aspen the only dog to complete all stages of Experiment 2. For Aspen, performance remained stable and above chance at onset asynchronies between 700 and 1,500 ms and dropped to chance levels at an onset asynchrony of 600 ms and below (Figure 5). The inflection point for sensitivity of the fitted function occurred at 633 ms, indicating that the discrimination threshold for odorant onset asynchrony lies in the range of 600-700 ms. However, the initial testing with larger increments for onset asynchrony suggested that onset asynchrony thresholds may be as low as 500 ms. Following completion of the study, three non-odour control sessions were completed, with a combined *d’* of -0.19, indicating no discrimination sensitivty when odorants were not present.

**Figure 5.**
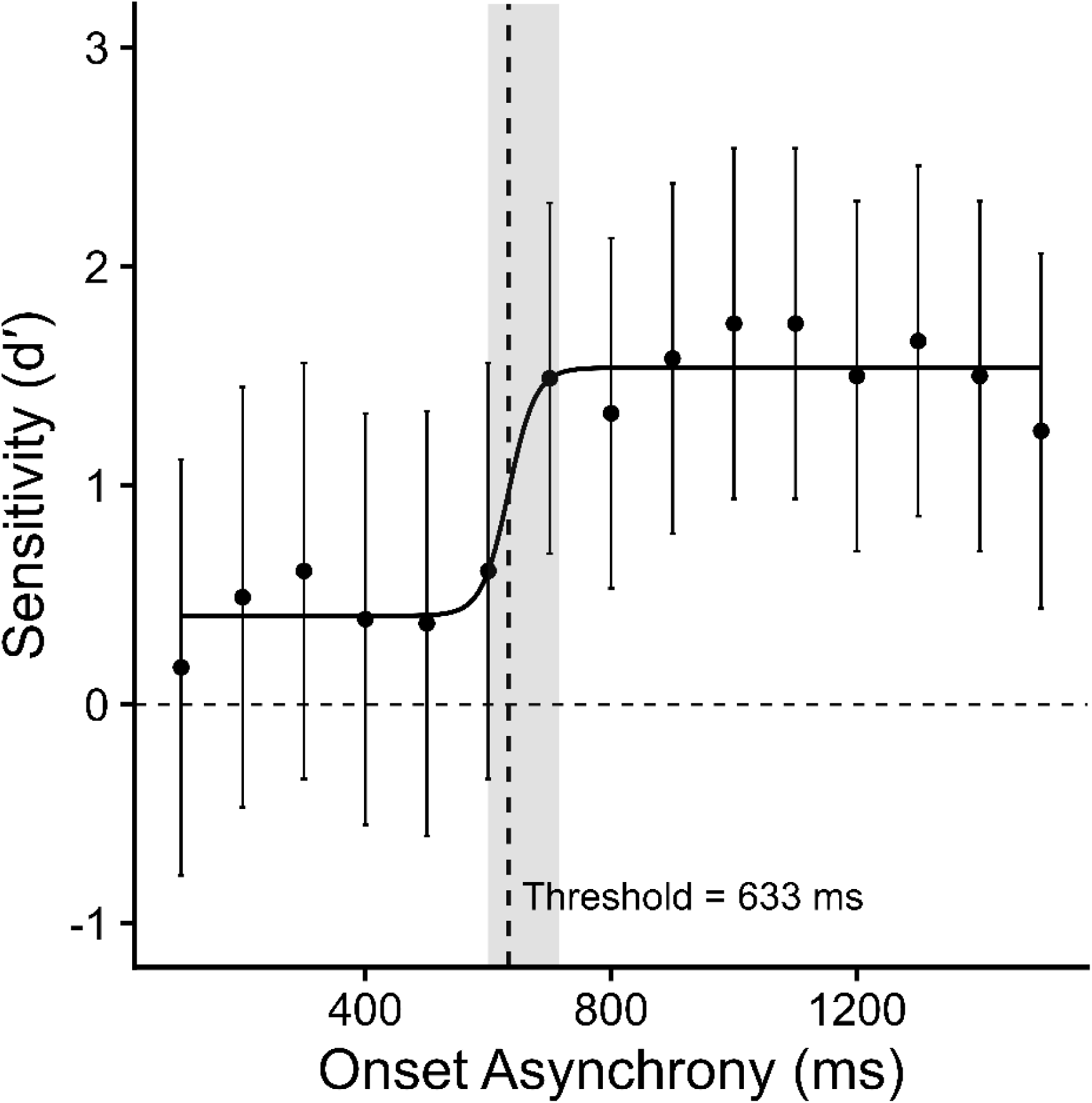
Discrimination between synchronous and asynchronous odorant mixtures with an onset asynchrony threshold of ∼633 ms. Error bars represent 95% confidence intervals around the calculated d’ value. Grey ribbon represents 95% confidence interval in the discrimination threshold estimate.

## Discussion

We found that a domestic dog can discriminate between binary odorant mixtures that differ only in the onset asynchrony of odorants, with an estimated discrimination threshold of approximately 633 ms. To our knowledge, this provides the first evidence that dogs can use onset asynchrony of odor stimulation to solve an olfactory discrimination task. This result adds to a growing body of evidence demonstrating that animals across diverse taxa are sensitive to the temporal structure of odor stimuli and can use temporal odor cues for odor discrimination, source segregation, and navigation and can use temporal odor cues for odor discrimination, source segregation, and navigation (Ackels, 2022; Marin et al., 2021; Szyszka et al., 2023).

Compared to insects, which can detect onset asynchronies as low as few milliseconds (Szyszka et al., 2023), vertebrates appear to require larger asynchronies to detect onset asynchrony. Mice can discriminate between synchronous and asynchronous odorant mixtures with onset asynchronies as low as 25 ms (Ackels et al., 2021), whereas research with humans has produced somewhat higher odorant onset asynchrony threshold estimates, ranging from 92 ms to 450 ms (Ding et al., 2019; Laing et al., 1994; Perl et al., 2020; Wu et al., 2024).

However, those previous studies with vertebrates used stimulus conditions that are not representative of naturally occurring onset asynchronies in odor plumes, as odorants were either presented in clean non-overlapping pulses (Ackels et al., 2021), or as single pulses coupled to the sniff phase (Wu et al, 2024; Perl et al., 2020; Ding et al., 2019; Laing et al., 1994). In human studies, odorant delivery was tightly coupled to the sniff cycle, by either presenting odorants directly into the nose or a short distance from the nostrils (Ding et al., 2019; Laing et al., 1994; Perl et al., 2020; Wu et al., 2024). In contrast, dogs in our study, encountered the odor stimuli repeatedly and at different times relative to their sniff cycle. This sampling mode more closely reflects natural conditions, in which animals dynamically interact with odor plumes while moving and may encounter odor stimuli at any point in their sniff cycle.

In the vertebrate study most similar to ours, Ackels et al. (2021) examined mice’s discrimination between synchronous (AB) and asynchronous (A-B) odorant mixtures, in which the asynchronous condition comprised sequential, non-overlapping stimuli of the two odorants within pulse trains. Under these conditions, mice discriminated between synchronous and asynchronous stimuli at onset asynchronies as low as 25 ms. In contrast, the asynchronous stimuli in our study involved substantial overlaps between the two odorants, with up to 58% overlap compared to 100% overlap in the synchronous condition. Thus, whereas Ackels et al. contrasted mixtures with 100% overlap (AB) and no overlap (A–B), our task required discrimination between two overlapping stimuli (AB vs A-AB-B). In addition, the asynchronous stimulus in our study comprised a sequence of pure and overlapping components: A alone, the AB mixture, and B alone. This increased compositional complexity likely made discrimination between synchronous and asynchronous mixtures more difficult, as discrimination could not rely on two simple sequences of two distinct odorants, but instead required interpretation of a temporally evolving stimulus that contained both pure and mixed components. This is an important distinction for ecologically relevant research because natural odor stimuli are unlikely to occur as fully non-overlapping (Celani et al., 2014).

Another key difference in the structure of odorant delivery in our study and that of Ackels et al. (2021) is that, in their paradigm, synchronous pulse trains included periods without odorants, whereas asynchronous pulse trains provided continuous olfactory stimulation. Under these conditions, animals could potentially discriminate between pulse trains based on differences in overall odorant presence or intensity, rather than on the onset asynchrony between odorants. In contrast, our stimuli included periods without odorants in both synchronous and asynchronous conditions, reducing such cues and increasing the requirement for discrimination based on onset asynchrony alone. This raises the possibility that, under comparable conditions, rodents may exhibit higher onset asynchrony thresholds than previously reported.

Given the methodological differences between our study, which prioritized complex odor sequences and more natural sampling, and prior research with vertebrates, meaningful comparisons between species are difficult. Although some differences may reflect anatomical or perceptual limits of the species tested, the current evidence is insufficient to support such conclusions.

A serendipitous outcome of this study was the demonstration of dogs’ apparent sensitivity to very small onset asynchronies, at least as low as 30 ms in auditory stimuli, which has not been studied previously with dogs. Following stimulus control tests, we identified that dogs were initially using subtle auditory cues generated by valve actuation to solve the task. Only after the development of a complex multi-valve masking system that effectively disguised these cues did behavior come under reliable olfactory control. This initial dominance of auditory cues highlights dogs’ sensitivity to multimodal stimuli in their environment, and it suggests that when olfactory cues are weak or ambiguous, dogs readily exploit alternative or more salient predictive signals. Moreover, this finding emphasizes the importance of rigorous stimulus control in psychophysics investigations.

The present findings also have important implications for understanding odor source segregation in dogs. Odorant onset asynchrony has been proposed as a general mechanism by which animals segregate odor sources in turbulent plumes (Hopfield, 1991). The ability of a dog to use odorant onset asynchrony supports this hypothesis, but the high onset asynchrony detection threshold observed here could suggest that dogs may rely more on other odor plume features of odor plumes. For example, dogs are highly sensitive to the concentration and relative strength of odorants, as demonstrated by generalization studies between training odors and higher concentration probes (Dechant et al., 2021), which may also facilitate the segregation of odor sources using concentration differences resulting from diffusion over several sniff cycles.

In addition to the sensory findings of this study, we developed an apparatus suitable for canine experiments that enables precise control of odorant onset asynchrony with low latency in odorant onset time and effective auditory masking. This apparatus may be useful for addressing related questions about temporal sensitivity in olfaction in future studies.

Although both the titration procedure (Experiment 1) and the method of constant stimuli (Experiment 2) had strengths and limitations, the method of constant stimuli appears to be a more efficient approach to establishing thresholds for this type of evaluation and may be beneficial to maintain participant engagement. Notably, as we found in this study, the titration procedure had a high rate of withdrawal and required a high number of sessions due to requiring large steps and multiple repetitions to account for learning effects over the course of the study. In comparison, with the method of constant stimuli, once dogs were trained, they only needed to complete a set number of sessions which were pre-defined before an estimate was made. This means that dog owners can be made aware of the time required at the start of the study and are less likely to have unexpected changes in circumstances.

However, the titration procedure is likely to give more accurate estimates for thresholds due to the ability to use smaller step sizes as the procedure continues. This can be achieved with the method of constant stimuli, but the number of trials needed to accommodate small enough onset asynchrony step sizes would result in the efficiency benefits being less prevalent.

Attrition due to loss of motivation reduced the final sample size, with only one dog completing the full experimental protocol. Replication with a larger cohort will be necessary to refine estimates of odorant onset asynchrony thresholds and determine the extent of inter-individual variability. In addition, as sniffing behavior was not measured concurrently, future studies integrating sniff-cycle monitoring with temporal discrimination tasks could provide insight into how active sampling dynamics interact with processing of temporal odor cues.

Finally, laboratory-generated odor pulses cannot fully capture the complexity of natural turbulent plumes, which fluctuate in odor concentration, pulse duration, intermittency, and odorant composition (Celani et al., 2014; Conchou et al., 2019; Gorur-Shandilya et al., 2019). Extending these methods to more naturalistic odor environments will therefore be an important next step in characterizing temporal olfactory sensitivity in canine olfaction.

## Data Availability

Data available on request.

